# Label-free quantification LC-mass spectrometry proteomic analysis of blood plasma in healthy dogs

**DOI:** 10.1101/2023.06.13.544743

**Authors:** Pavlos G. Doulidis, Benno Kuropka, Carolina Frizzo Ramos, Alexandro Rodríguez-Rojas, Iwan A. Burgener

## Abstract

Bloodwork is a widely used diagnostic tool in veterinary medicine, as diagnosis and therapeutic interventions often rely on blood biomarkers. However, biomarkers available in veterinary medicine often lack sensitivity or specificity. Mass spectrometry (MS)-based proteomics technology has been extensively used in biological fluids and offers excellent potential for a more comprehensive characterization of the plasma proteome in veterinary medicine. In this study, we aimed to identify and quantify plasma proteins in a cohort of healthy dogs and compare two techniques for depleting high-abundance plasma proteins to enable the detection of lower-abundance proteins. We utilized surplus lithium-heparin plasma from 30 healthy dogs, which were subdivided into five groups of pooled plasma from 6 randomly selected individuals each. Our goal was to identify and quantify plasma proteins via label-free quantification LC-mass spectrometry. Additionally, we employed different methods to deplete the most abundant proteins. Firstly, we used a commercial kit for the depletion of high-abundance plasma proteins. Secondly, we employed an in-house method to remove albumin using Blue-Sepharose. Among all the samples, some of the most abundant proteins identified were apolipoprotein A and B, albumin, alpha-2-macroglobulin, fibrinogen beta chain, fibronectin, complement C3, serotransferrin, and coagulation Factor V. However, neither of the depletion techniques achieved significant depletion of high-abundant proteins. Nevertheless, the two different depletion methods exhibited substantial differences in the fold-change of many proteins, suggesting partial depletion that did not contribute to an increase in the number of detected proteins. Despite this limitation, we were able to detect and quantify many clinically relevant proteins. The determination of the healthy canine proteome is a crucial first step in establishing a reference proteome for canine plasma. This reference proteome can later be utilized to identify protein markers associated with different diseases, thereby contributing to the diagnosis and prognosis of various pathologies.

## Introduction

Biomarkers are defined as traits that can be measured as an indicator of a pathogenic process or a pharmacologic response to treatment and can be assessed objectively (Fidock and Desilva, 2012). Enzymatic assays and immunoassays are the most commonly used methods for quantifying highly abundant biomarkers. However, the rate of introducing novel biomarkers in medicine is reported to be less than two per year (Anderson et al., 2013), and the adoption of these biomarkers in veterinary medicine is often delayed.

Mass spectrometry (MS)-based proteomics has emerged as a powerful technology in biological (Ahrens et al., 2010; Cox and Mann, 2011; Bensimon et al., 2012) and medical (Huang et al., 2017; Murphy et al., 2018) research. It offers the capability to comprehensively characterize the plasma proteome, thereby contributing to the discovery of new biomarkers (Ndao, 2012; Aebersold and Mann, 2016; Geyer et al., 2017). Unlike traditional techniques, mass spectrometry provides high-precision peptide masses and fragmentation spectra derived from sequence-specific digestion of proteins of interest (Geyer et al., 2017). Proteomics is highly specific due to the uniqueness of peptide masses and sequences, in contrast to colorimetric enzyme tests and immunoassays (Wild & Davies, 2013).

Among various approaches, liquid chromatography-tandem mass spectrometry (LC-MS/MS) is the preferred method in medical research due to its high analytical specificity and sensitivity, enabling the detection and quantification of low-abundance proteins, such as drugs and metabolites (Vogeser and Parhofer, 2007). Consequently, MS-based proteomic techniques are gaining increasing interest in small animal veterinary medicine. Although research on MS-based proteomic analysis in veterinary medicine is not as extensive as in human clinical research, studies analyzing biological fluids of dogs with various diseases have been reported (Adaszek et al., 2014a; Escribano et al., 2016a; Locatelli et al., 2016; Martinez-Subiela et al., 2017a; Franco-Martínez et al., 2020; Lucena et al., 2020; Phochantachinda et al., 2021). Given the limited technical and medical data in the veterinary literature and the importance of proteomic analysis for detecting novel biomarkers or biomarker networks in canine plasma, there is a need to explore new methods for studying blood biomarkers.

Plasma refers to the liquid portion of uncoagulated blood left behind after all cell types are removed. Heparin is one of the most commonly used anticoagulants for plasma preparation, acting through anti-thrombin activation. When clot formation in plasma is triggered to form serum, fibrinogen and other coagulation factors are depleted, while activation peptides like fibrinopeptide A and B are released (Brummel et al., 1999; Profumo et al., 2003; Banks et al., 2005). Over the last decades, the number of proteins identified in human plasma has increased exponentially (Anderson et al., 2004; Omenn et al., 2005). Plasma proteomic analysis offers the advantage of quantifying high-abundance proteins such as fibrinogen and partially other coagulation factors that are no longer present in serum, potentially revealing specific patterns in diseases. However, due to plasma’s extremely high dynamic range, identifying lower abundance proteins by LC-MS can be challenging. Techniques like antibody depletion of abundant plasma proteins such as albumin and fibrinogen, as well as extensive plasma fractionation, have been successfully used to facilitate the detection of low-abundance proteins in human samples (Tu et al., 2010; Bellei et al., 2011; Cao et al., 2012). These techniques have also been combined successfully to identify several thousand proteins (Liu et al., 2006; Pan et al., 2011; Keshishian et al., 2015).

The aim of this study is to utilize label-free quantification LC-MS proteomic analysis to explore the composition of canine plasma in healthy dogs from different breeds. These findings will serve as a foundation for future studies involving larger disease-specific cohorts, with the goal of detecting new biomarkers or gaining a better understanding of biomarker networks in various diseases. Additionally, we aim to contribute to the establishment of a reference database for dogs by depositing the list of detected proteins to PeptideAtlas (Schwenk et al., 2017).

## Results and Discussion

Two-dimensional gel electrophoresis (2-DE) has been widely used for proteomic analysis (Herosimczyk et al., 2006; Desrosiers et al., 2007). However, this method has limitations in effectively identifying proteins in low abundance and its dynamic range is limited. Gel-free methodologies have gained attention in recent years since they allow for the determination and quantification of a wider range of proteins (Marcus et al., 2020). The need for detecting novel prognostic biomarkers to predict disease outcomes has led proteomic research in canine medicine to focus on infectious diseases, with leishmaniosis being the most prominent example (Agallou et al., 2016; Escribano et al., 2016b; Martinez-Subiela et al., 2017b; Franco-Martínez et al., 2019), followed by diseases like babesiosis (Adaszek et al., 2014b; Galán et al., 2018; Winiarczyk et al., 2019), dirofilariasis (Hormaeche et al., 2014), ehrlichiosis (Escribano et al., 2017), and parvovirus infection (Franco-Martínez et al., 2018). Some studies on patients with leishmaniosis and babesiosis found a significant downregulation of Apolipoprotein A, which may reduce the individual’s capacity to respond to oxidative damage. Beyond the field of infectious diseases, proteomic analysis has provided new insights in veterinary nephrology, revealing that an increase in proteins like retinol-binding protein predicts kidney damage before azotemia develops (Nabity et al., 2011; Chacar et al., 2017; Ferlizza et al., 2020), and in veterinary endocrinology, uncovering the role of Apolipoprotein I in canine obesity (Tvarijonaviciute et al., 2012; Lucena et al., 2019). According to the literature, the most commonly analyzed samples in veterinary medicine are serum and saliva (González-Arostegui et al., 2022). Studies have also been conducted with other biological fluids such as cerebrospinal fluid, bile, liver, synovial fluid, myocardium (Yuan et al., 2006; Kjelgaard-Hansen et al., 2007; Plumb et al., 2009; Nakamura et al., 2012; Lawrence et al., 2018), and feces (Cerquetella et al., 2019). Currently, there is no systematic catalog of dog plasma proteins available. However, a catalog is available for human plasma that can be used for comparison, such as PeptideAtlas, which contains 3509 proteins (Schwenk et al., 2017). Although proteomic analysis of canine plasma is less common than serum analysis, some studies on canine plasma have been reported (Kuleš et al., 2016; Tvarijonaviciute et al., 2016; Phochantachinda et al., 2021).

In our study, we utilized a label-free quantification LC-MS method to analyze canine plasma. The plasma of 30 healthy individuals was used. The median age of the 30 dogs was 5,7 years old (range 1,3-9,9 years old), median weight was 18,4 kg (range 5-37,5 kg), while 15 dogs were male (50%) and 15 dogs were female (50%). Sixteen (60%) dogs were castrated and 14 (40%) were intact. All included dogs had a body condition score of 5/9. Of these dogs, 9 were mix breeds, 3 Australian Shepherds, 2 Golden Retrievers, 2 German Shepherds, 2 Puli’s and 12 other breeds were represented by 1 dog each. These samples were randomly selected, according to the inclusion criteria, to avoid preserve heterogenicity and were thereafter grouped into five pools of six individuals using a computer script, as outlined in the material and methods section. We assessed the plasma protein content and evaluated two depletion methods for high-abundance proteins. The first method involved the use of a commercial kit (referred to as “kit”), while the second method employed a low-cost in-house approach using Blue-Sepharose. The purpose of the Blue-Sepharose method was specifically to remove albumin (referred to as “Blue-Sepharose”). To optimize the process, we incorporated a set of clinical samples obtained from healthy individuals with well-established normal profiles of routine blood parameters.

All raw data and result files from our MS experiments have been made available in the public repository ProteomeXchange, as outlined in the materials and methods section. Initially, we identified a total of 282 proteins. Subsequently, proteins that were not identified in all three replicates from at least one of the three groups (Control, Kit, Blue-Sepharose) were filtered out from the quantification. This filtering process resulted in the quantification of 181 proteins in the plasma samples. Figure 1 illustrates the above-mentioned data processing steps. For protein description, gene IDs were primarily used. In cases where gene names were unavailable, protein IDs were utilized instead. Among all the samples, some of the most abundant proteins identified were apolipoprotein A and B (APOA, APOB), albumin (ALB), alpha-2-macroglobulin (A2M), fibrinogen beta chain (FGB), fibronectin (FN1), complement C3 (C3), serotransferrin (LOC477072), coagulation Factor V (F5), maltase-glucoamylase (MGAM), and several uncharacterized proteins (LOC611458, LOC481722, A0A8I3P3U9). For a detailed list of all identified proteins and the raw data, please refer to the Supplementary Material (Supplementary Table S1).

**Figure 1.**
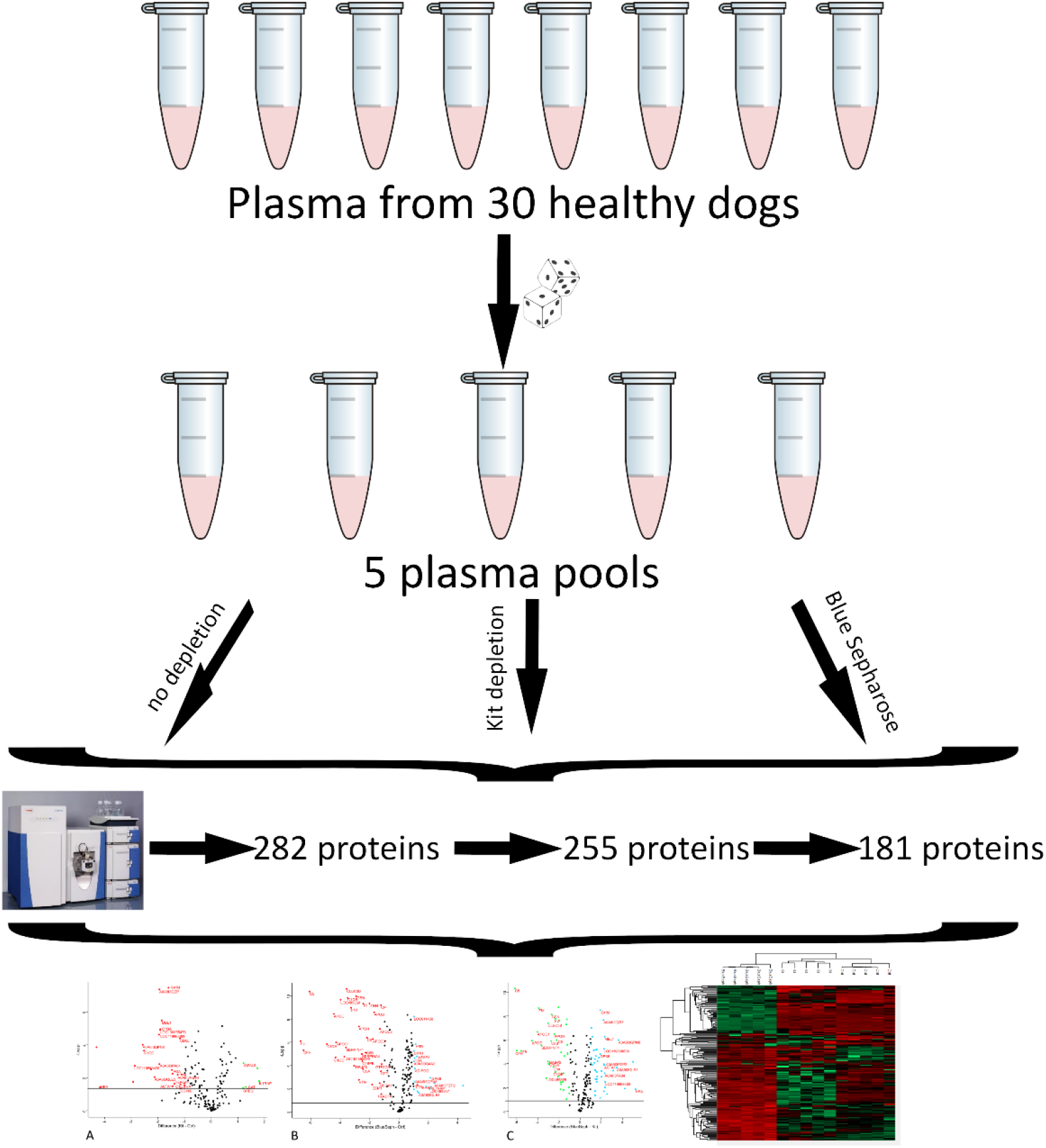
Illustration of the experimental setup and data processing for the canine plasma proteome analysis used in this study. The pools five pools were generated using a computer script to ensure the randomization of each sample. Initially 282 proteins were detected. After removing reverse hits and contaminants 255 proteins remained and after excluded proteins not identified in all 3 replicates of at least one experiment a total of 181 proteins was analysed.

Prior to excluding proteins that were not detected in at least three replicates from any of the three groups, 163 out of the 192 proteins were present in all three groups, while 15 proteins were assigned to the Control and Kit groups, 5 proteins to the Control and Blue-Sepharose groups, and 5 proteins to the Kit and Blue-Sepharose groups. Additionally, one protein (S100A12) was exclusively identified in the kit depletion experiment, two proteins (Ig-like domain-containing proteins, Protein IDs: A0A8I3P3T7 and A0A8I3P941) were exclusively identified in the Blue-Sepharose depletion experiment, and one protein (AMBP) was exclusively identified in the experiment without depletion (control).

The ranking of protein abundance can be observed in Figure 2. Principal component analysis demonstrated significant and clear segregation among the three methods (Figure 3). The pairwise hierarchical clustering and correlation analysis of all protein samples using the two different depletion methods, as well as the total proteins detected without depletion as control, are depicted in Figure 4. Identifying low-abundance proteins in plasma poses a challenge due to its wide dynamic range.

**Figure 2.**
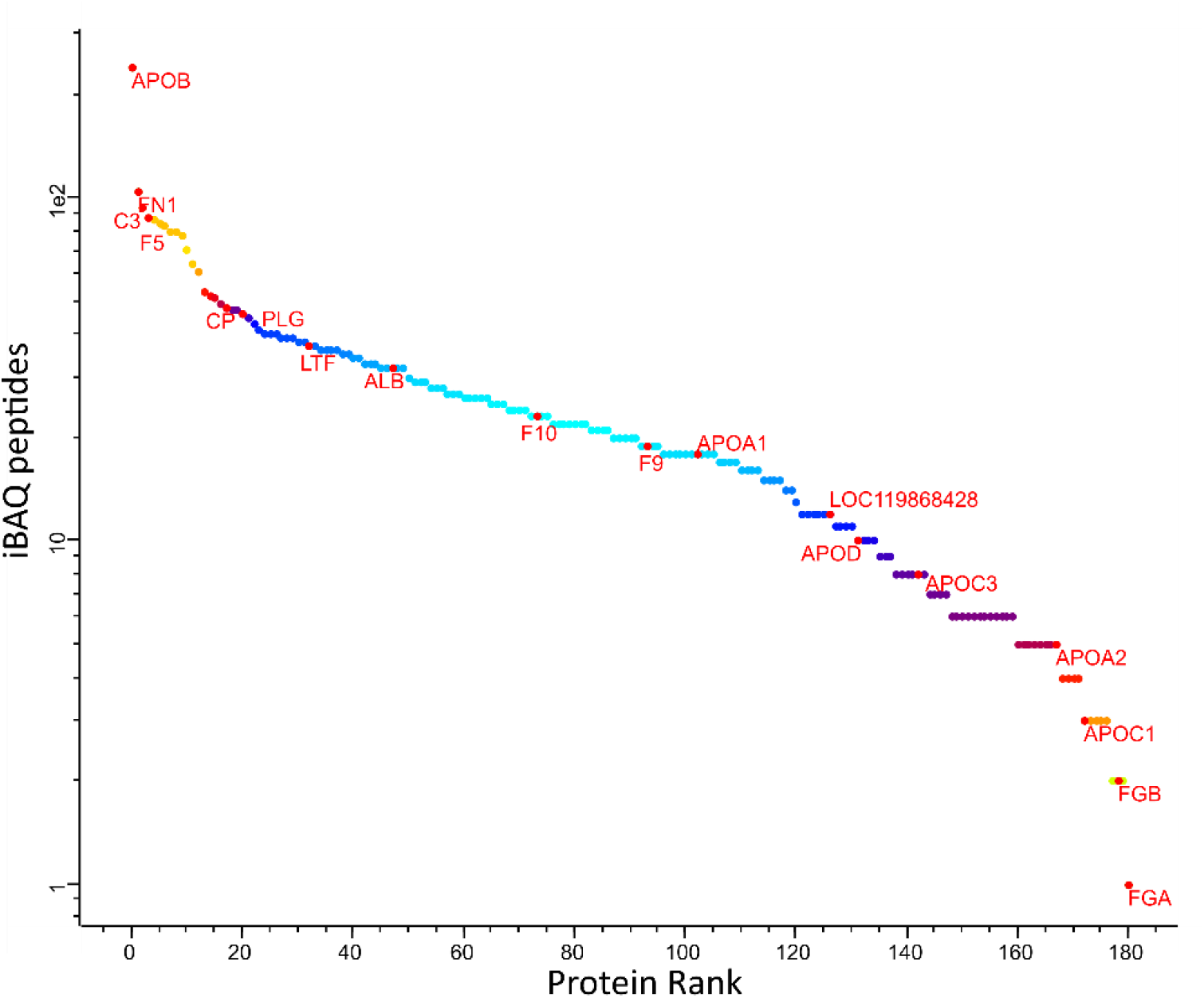
Abundance rank of the identified canine plasma proteins. Selected protein representing different level of abundances are labelled: APOB: Apolipoprotein B, FN1 Fibronectin 1, C3: Compliment 3, F5: Coagulation Factor 5, CP: Ceruloplasmin, PLG: Plasminogen, LTF: Lactoferrin, ALB: Albumin, F10: Coagulation Factor 10, F9: Coagulation Factor 9, APOA1: Apolipoprotein A1, LOC119868428: Ferritin, APOD: Apolipoprotein D, APOC3: Apolipoprotein C3, APOA2: Apolipoprotein A2, APOC1: Apolipoprotein C1, FGB: Fibrinogen beta chain, FGA: Fibrinogen alpha chain.

**Figure 3.**
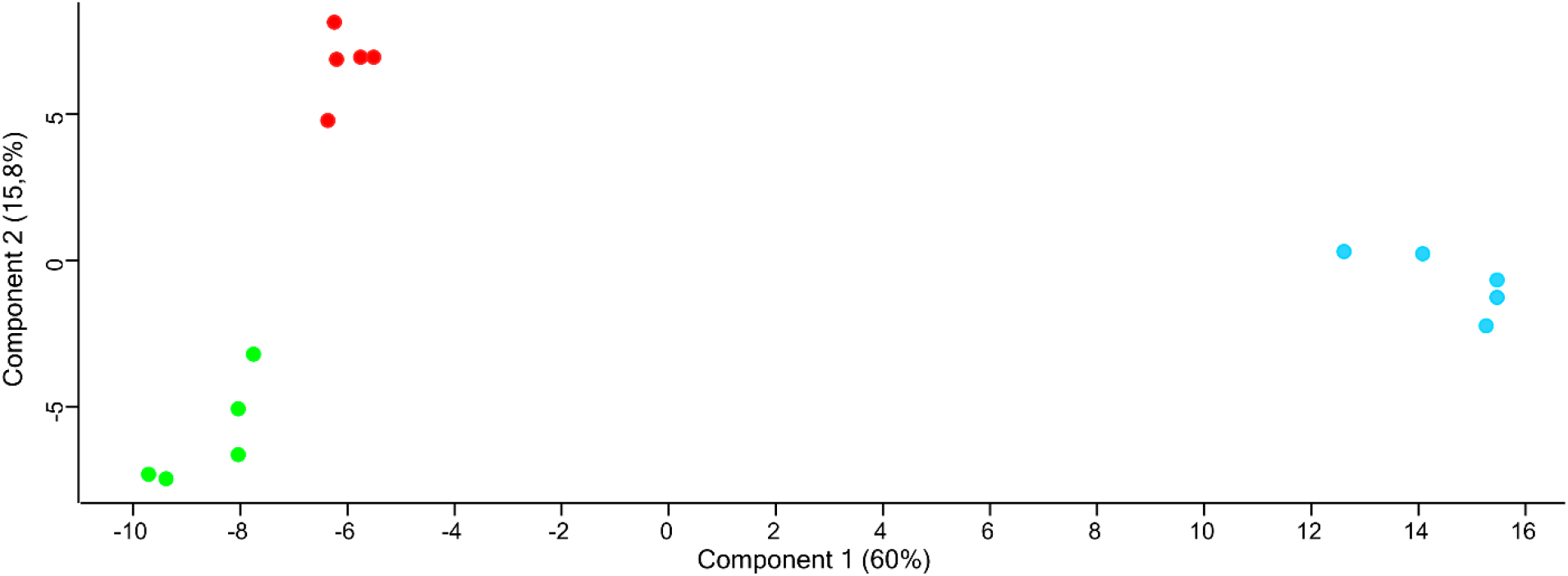
Principal Component Analysis of the three different methods used. A clear segregation can be seen between the three groups. (Control in red, Kit in green, and Blue-Sepharose in light blue).

**Figure 4.**
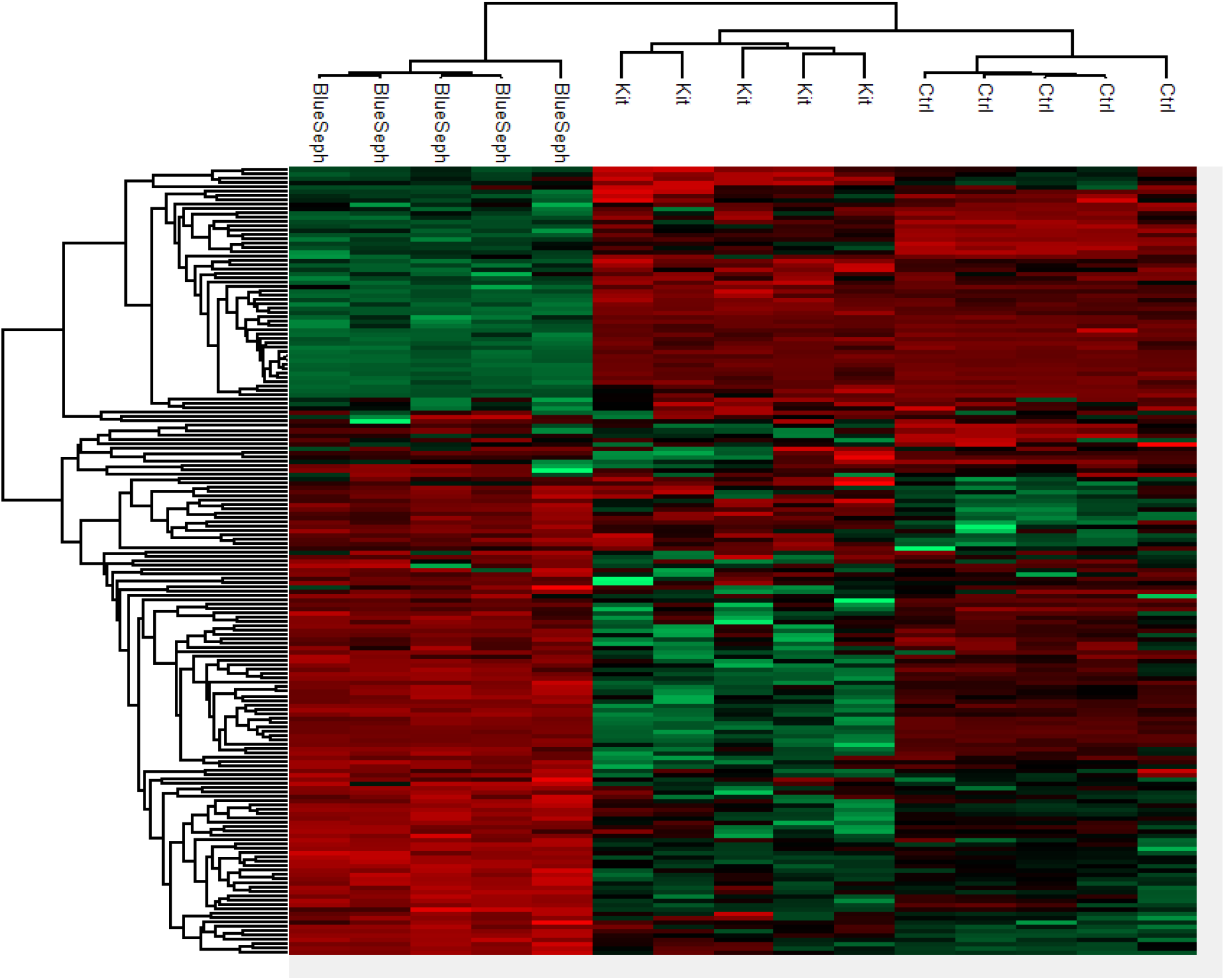
Heatmap of the detected proteins, presenting the result of a two-way hierarchical clustering of the proteins found in the three groups. The diagram was constructed using the complete-linkage method together with the Euclidean distance. Each row represents a differentially abundant proteins and the columns are the different samples tested (5 pools in each group). The intensity scale illustrates the relative level of differentially protein concentrations with green portraying up-represented and red down-represented.

Significant depletion of high-abundance proteins, particularly albumin, was not achieved with either technique (Kit, Blue-Sepharose) compared to the control technique without depletion (Supplementary Figure S1). However, the log2 fold-change of albumin concentration in samples after using the kit and the Blue-Sepharose depletion was small (0.312) but statistically significant (q value = 0.007), with albumin being more abundant after depletion with Blue-Sepharose. One possible explanation for the inadequate depletion of high-abundance canine plasma proteins with the commercial kit is that the kit used in this study is designed for depleting high-abundance proteins in humans and has not been validated for canine proteins. These antibodies are not entirely specific and have affinity for several other proteins (Bellei et al., 2011). Furthermore, the Blue-Sepharose depletion method used in this experiment is not yet standardized for animal samples, and we hypothesize that variations in the protocol, such as the amount of sample or Blue-Sepharose added, might contribute to better depletion results.

In this study, we used canine plasma instead of serum as we aimed to identify proteins involved in the coagulation cascade. Fibrinogen A (FGA) showed a significant decrease in abundance (log2 fold-change -1.469, q-value = 0.006) after depletion with the kit compared to the no-depletion group. However, it is still unclear to what extent fibrinogen affects depletion and how it interferes with the detection of lower abundance proteins.

The two different depletion methods exhibit significant differences in the fold change of numerous proteins compared to the three different techniques (Figure 5A, 5B, 5C). Thirty-two proteins were differentially abundant among the control and kit depletion methods, with 27 being more abundant with the control method and 5 with the kit depletion method (Supplementary Table S2). Among the most important proteins that showed a significant increase after kit depletion compared to the control method are interleukin 1 receptor accessory protein (IL1RAP), solute carrier family 12 member 4 (LCAT), insulin-like growth factor binding protein acid labile subunit (IGFALS), sex hormone-binding globulin (SHBG), and V-type proton ATPase subunit G (A0A8I3PF02). On the other hand, hemoglobin subunit alpha (HBA), ferritin (LOC119868428), complement C1q C (CIQC), fibrinogen alpha chain (FGA), and Ig-like domain-containing protein (A0A8I3PB96) were significantly more abundant in the control experiment without depletion (Figure 6A). Fifty-three proteins were differentially abundant among the control and Blue Sepharose depletion methods, with 35 being more abundant with the control method and 18 with the Blue Sepharose depletion method (Supplementary Table S3). Interestingly, IL1RAP and SHBG were quantified in significantly increased concentrations (fold change) in samples after depletion with the Blue-Sepharose method compared to the control method (Figure 6B). Finally, we compared the protein profiles detected with the two different depletion techniques (Figure 6C). Eighty-two proteins were differentially abundant among the Blue Sepharose and kit depletion methods, with 45 being more abundant with the Blue Sepharose method and 37 with the kit depletion method(Supplementary Table S4). The most noteworthy proteins that were significantly more abundant after kit depletion include complement C5 (C5), coagulation Factor V (F5), apolipoprotein E (APOE), fibronectin (FN1), and serpin family F member (SERPINF1), while HBA, Ig-like domain-containing protein (A0A8I3QPN8), ferritin (LOC119868428), C-type lectin domain-containing protein (MBL1), and immunoglobulin heavy constant mu (IGHM) were found in significantly higher concentrations using the Blue-Sepharose method.

**Figure 5.**
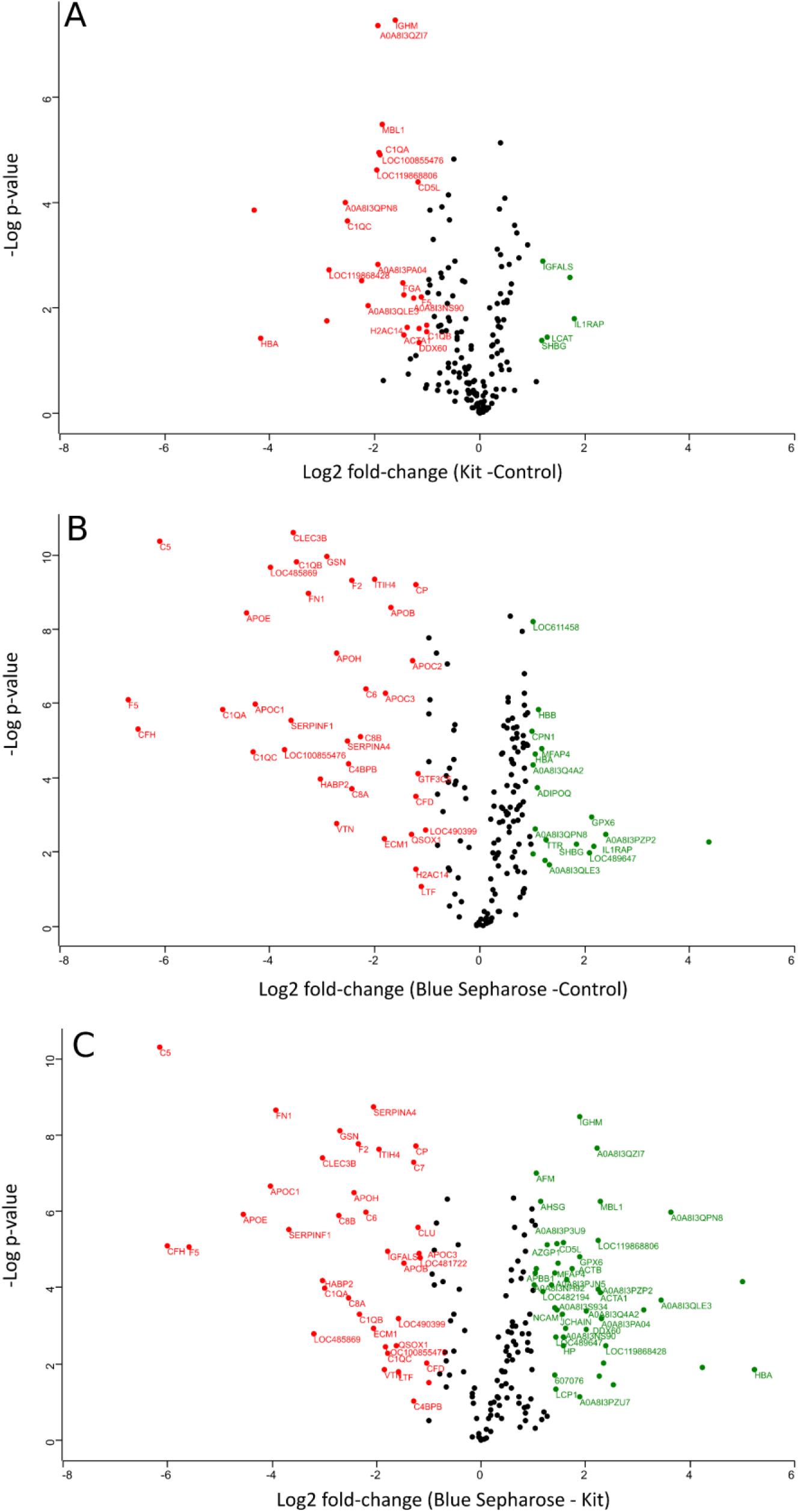
**A, B, C**. Volcano plots –log p values versus a log2-fold change of protein intensity measured by LC-MS of canine plasma proteins from the three conditions of plasma processing in this study. Black dots represent non-significant differentially abundant proteins, green dots show the up-represented fraction and, and red ones represent the down-represented fraction.

**Figure 6.**
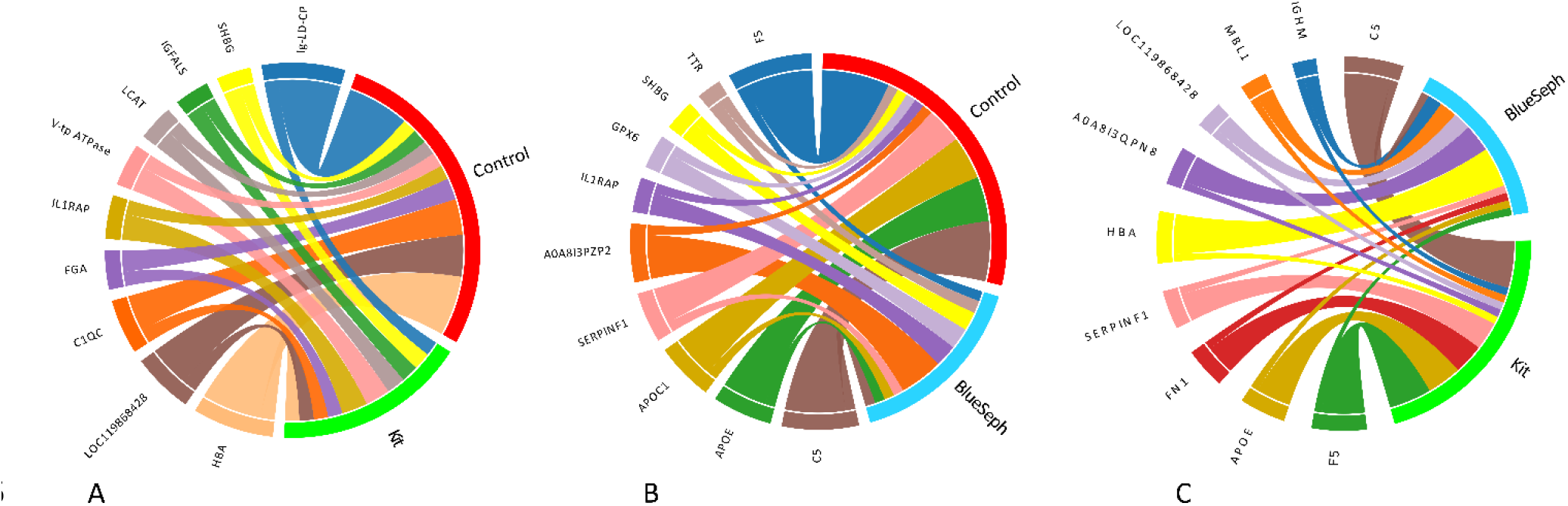
**A, B, C**. Chord diagrams showing the associations among differentially abundant proteins of Control, Kit, and Blue-Sepharose (Blue Seph) groups. Only the most important abundant proteins detected are shown. For better comprehension, each circle includes two comparisons each time (Control in red, Kit in green, and Blue-Sepharose in light blue). The chord diagrams show the key proteins identified by their comparative abundance. The outer ribbon identifies the respective groups-experiment and encompasses the perturbed protein quantification with each method. Chords connect proteins related to more than one method in the inner circle. Only significant hits are represented in these chord charts (at least q<0.05).

In our experimental conditions, we did not find a clear benefit of using depletion methods because one of the goals of using these methods was to increase the number of detected proteins, which was not achieved in our study. There is a possibility that the kit for protein depletion is not well optimized for dog plasma. This seems to be the case for Blue-Sepharose as well, as albumin did not significantly decrease and other unexpected proteins showed a decrease, likely due to nonspecific binding. For this reason, in the future, with the present instrumental setup, we consider that it is not necessary, in principle, to use a depletion method for canine plasma analysis, or alternatively, a new specific depletion method should be developed. In the case of Blue-Sepharose, the diminishing abundance of certain proteins, but not albumin, may indicate nonspecific binding of these protein sets. In contrast, albumin unexpectedly did not decrease in concentration.

## Conclusions

Label-free LC-mass spectrometry proteomic analysis can reliably detect and quantify multiple proteins in canine plasma, making it a valuable tool for unraveling the pathogenesis of various diseases in veterinary medicine. It provides influential information for accurate disease diagnosis and prognosis estimation in future analyses. Our results indicate that protein depletion with the two methods described herein is not adequately achieved. Although the abundance of various proteins can differ significantly, these methods, as performed here, do not contribute to the determination of lower abundance proteins in canine plasma. Prospective controlled studies in animal disease models are expected to shed light on the utility of gel-free label-free LC-mass spectrometry proteomic analysis in veterinary medicine.

## Material and methods

### Animals and sample collection

Clinically healthy client-owned dogs (N=30) with no signs of disease within the last two months were presented at the Division for Small Animal Internal Medicine (Veterinary University of Vienna, Vienna, Austria) over a period of six months for clinical examination and blood sampling. They were enrolled in a study (Ref: BMBWF 20221-0.210.26) conducted by the Division for Small Animal Internal Medicine and the Interuniversity Messerli Institute of Research (Veterinary University of Vienna, Austria). Prior to enrollment, written informed consent was obtained from the owners. Dogs of different breeds, body weights, and genders, ranging in age from 1 to 10 years old, were considered for the proteomics study. Inclusion criteria involved a comprehensive evaluation, including a detailed history, physical examination, and blood sampling performed by two authors (PGD, CFR). Complete blood count (CBC), serum biochemical profile, and electrolyte measurements were conducted. Additionally, the body condition score (Nestle Purina scale: ranging from 1-very thin to 9-significant obesity) of each dog was recorded. Dogs younger than one year, weighing less than 5kg, and those with any pathological signs or recent medication administration within the last two months were excluded from the study. Likewise, dogs with significant alterations in any blood parameters were not enrolled. Randomization and group selection were performed using the “tidyverse” R package.

### Quantitative proteome analysis by label-free liquid chromatography–mass spectrometry (LC–MS)

Blood samples were collected using 2 ml Vacuette tubes with lithium heparin 13×75 green cap-white ring PREMIUM (Greiner Bio-One GmbH, Bad Haller Str. 32, 4550, Austria). After centrifugation at 2000 x g for 5 minutes, the plasma was separated and stored at -21°C. Five microliters of plasma from each of the 30 individuals was collected by centrifugation. We created three separated groups that consisted of five pools, each containing pooled plasma from six individuals. One of the groups went for 14 more abundant protein depletion procedure using a commercial kit (Top14 Abundant Protein Depletion Mini Spin Columns, Thermo Scientifics, Germany) that was used following the instructions of the manufacturer. The second group consisted of in-house albumin depletion procedure using blue sepharose CL6B (GE Health Care, Germany). The procedure consisted in mixing 50 μl of plasma pools with 100 μl of pre-equilibrated blue sepharose with phosphate buffer (150 mM, pH 7.2), incubation for 30 minutes with soft shaking, separation of the supernatant by centrifugation (4000 x g for 10 minutes), and separation of the left over that was used for the proteomic procedure. The third group consisted in pools of plasma without any treatment that were used directly for proteomics.

Five microliters of plasma per pool was transferred to a tube containing 20 μl of urea denaturing buffer (6 M urea, 2 M thiourea, and 10 mM HEPES; pH 8.0). Disulfide bonds from the plasma proteins were reduced by adding 1 μl of dithiothreitol (10 mM) and incubated for 30 minutes at room temperature. Afterwards, the samples were alkylated by adding 1 μl of iodoacetamide (55 mM) solution and incubated at room temperature for another 30 minutes in the dark. The samples were diluted with four volumes of ammonium bicarbonate buffer (40 mM) and digested overnight at 37°C by adding 1 μl of trypsin protease (Thermo Scientific, USA) (1 μg/μl). To acidify the samples, 5% acetonitrile and 0.3% trifluoroacetic acid (TFA; final concentration) were added, and subsequently, the samples were desalted using C18 StageTips with Empore™ C18 Extraction Disks (Rappsilber et al., 2007). The peptides eluted from the StageTips were dried using vacuum centrifugation.

Peptides were reconstituted in 20 μl of a solution containing 0.05% TFA and 4% acetonitrile. Then, 1 μl of each sample was applied to an Ultimate 3000 reversed-phase capillary nano liquid chromatography system connected to a Q Exactive HF mass spectrometer (Thermo Fisher Scientific). The samples were injected and concentrated on a PepMap100 C18 trap column (3 μm, 100 Å, 75 μm inner diameter [i.d.] × 20mm, nanoViper; Thermo Scientific) that was equilibrated with 0.05% TFA in water. After switching the trap column inline, LC separations were performed on an Acclaim PepMap100 C18 capillary column (2 μm, 100 Å, 75 μm i.d. × 250mm, nanoViper; Thermo Scientific) at an eluent flow rate of 300 nl/min. Mobile phase A consisted of 0.1% (v/v) formic acid in water, while mobile phase B contained 0.1% (v/v) formic acid and 80% (v/v) acetonitrile in water. The column was pre-equilibrated with 5% mobile phase B, followed by an increase to 44% mobile phase B over 100 minutes. Mass spectra were acquired in a data-dependent mode, utilizing a single MS survey scan (m/z 350–1650) with a resolution of 60,000, and MS/MS scans of the 15 most intense precursor ions with a resolution of 15,000. The dynamic exclusion time was set to 20 seconds, and the automatic gain control was set to 3 × 10^6^ and 1 × 10^5^ for MS and MS/MS scans, respectively.

MS and MS/MS raw data analysis was performed using the MaxQuant software package (version 2.0.3.0) with the implemented Andromeda peptide search engine (Tyanova et al., 2016a). The data were searched against the *Canis lupus familiaris* reference proteome (ID: UP000002254; downloaded from Uniprot.org on 10.1.2022; 43,621 sequences) using the default parameters and enabling the options of label-free quantification (LFQ) and match between runs. Data filtering and statistical analysis were conducted using the Perseus 1.6.14 software (Tyanova et al., 2016b). Only proteins that were identified and quantified with LFQ intensity values in at least three (out of five) replicates within at least one of the three experimental groups were used for downstream analysis. Missing values were replaced from a normal distribution (imputation) using the default settings (width 0.3, downshift 1.8). Mean log2-fold differences between groups were calculated in Perseus using Student’s t-test. Proteins with a minimum 2-fold intensity change compared to the control (log2-fold change ≥ 1 or log2-fold change ≤ -1) and a p-value ≤ 0.05 were considered significantly abundant.

## Statistical analysis

All statistical comparisons between groups were performed using student’s t-Test implemented by the Perseus computational platform. Adjusted *P*-values after FDR (*q*-values) were considered significant for values below 0.05.

## Acknowledgments

We are also grateful to the mass spectrometry unit I of the Core Facility BioSupraMol of the Free University of Berlin, Germany, which is supported by the DFG.

## Legend to the figures and tables

**S1 Figure**. Venn diagram of differentially abundant proteins of plasma from Control, Kit, and Blue-Sepharose groups.

**S1 Table**. Output table of the proteomic experiment reporting plasma protein detection and quantification from dog plasma. The data includes treating three conditions: total plasma, depletion fraction (Thermo Scientific depletion kit), and in-house albumin depletion using Blue-Sepharose. The same procedure was applied to five dog plasma pools (six individuals each). Statistical analysis used student t-test and false discovery rate (FDR) to correct the p-values (data analysis using MaxQuant and Perseus software for label-free quantification of proteins with LC-MS).

**S2 Table**. Table showing the differentially abundant proteins among Control and Kit methods including protein ID, Gene name, fold-change and statistical significance.

**S3 Table**. Table showing the differentially abundant proteins among Control and Blue Sepharose methods including protein ID, Gene name, fold-change and statistical significance.

**S4 Table**. Table showing the differentially abundant proteins among Blue Sepharose and Kit methods including protein ID, Gene name, fold-change and statistical significance.

## Notes

### Competing Interest Statement

The authors have declared no competing interest.

