## Supplementary figures and images for "Label-free quantification LC-mass spectrometry proteomic analysis of blood plasma in healthy dogs"

### Supplementary Figure S1

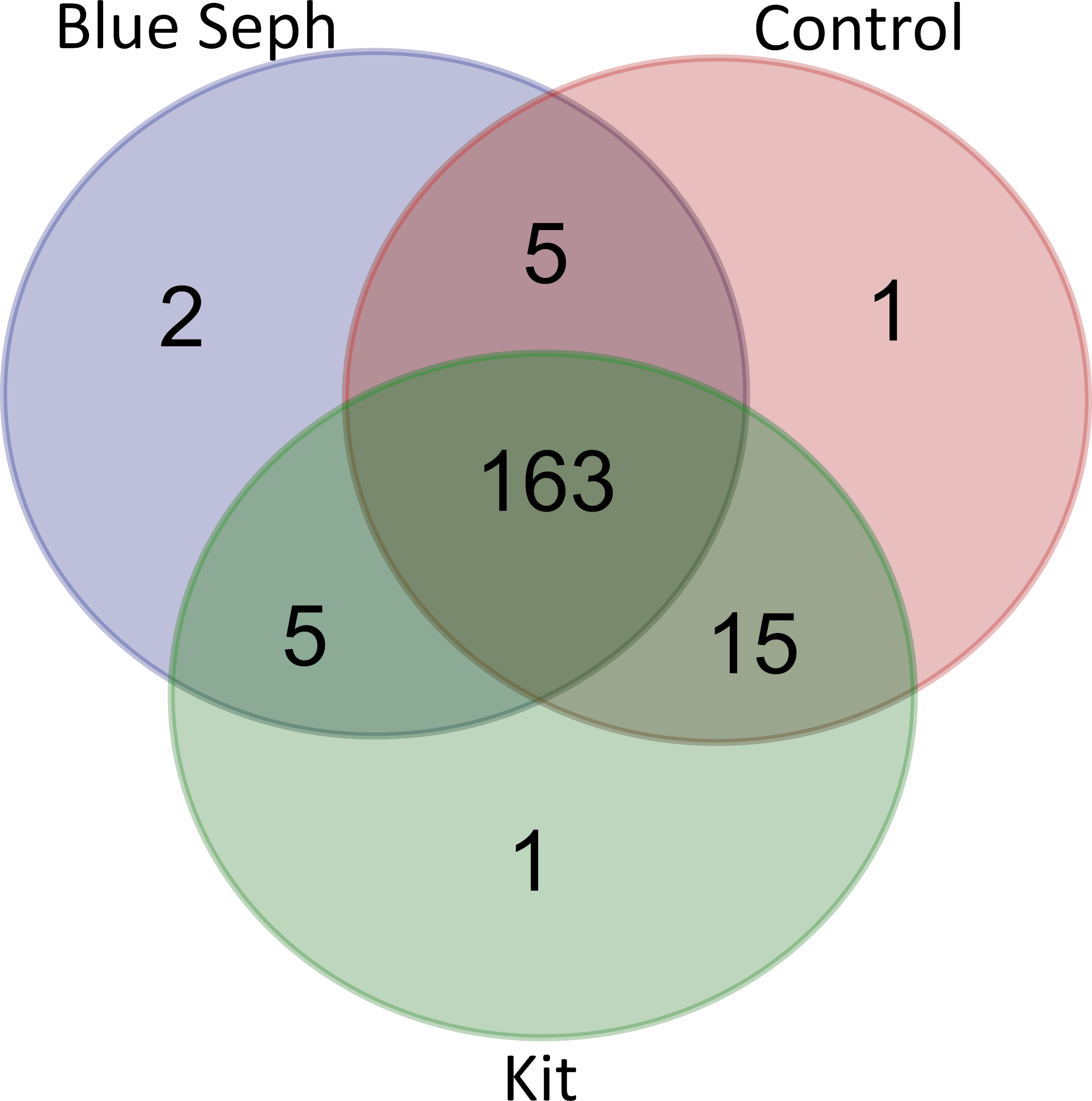
